# The Mechanical of Organic Acids Secreted by Roots of Tartary Buckwheat under the Effects of Low Nitrogen Stress

**DOI:** 10.1101/346064

**Authors:** Wei Chen, Yaru Cui, Yang Yang, Qianhua Huangfu, Congjian Sun

## Abstract

A pot experiment was conducted to study the effects of two different low nitrogen tolerant tartary buckwheat varieties’ (Diqing buckwheat (DQ, low nitrogen resistance) and Heifeng 1 (HF, low nitrogen sensitive) response mechanism of organic acids to low nitrogen stress. The results showed that the soil moisture of HF and DQ under low nitrogen treatment decreased 24.2% and 14.32%, respectively when compared with normal nitrogen treatment, and the water consumption of DQ was significantly higher than that of HF at seedling stage. Under low nitrogen treatment, the soil pH value of DQ was 1.44% and 8.44% lower than that of HF at seedling and flowering stages, respectively, the content of NH_4_^+^ in DQ soil was 8.2% lower than that of HF at maturity stage, the content of NO_3_^−^ was significantly higher than that HF 49.2%, 12.9%, and 16.6% in each growth period, respectively. Split plot analysis showed that nitrogen treatment significantly affected the organic acids content in the soil of the buckwheat. The secretion content of organic acids are different among buckwheat cultivars under low nitrogen stress. In the soil of DQ, the content of malonic acid was higher than that of HF by 34.39% at maturity stage; the content of oxalic acid was respectively higher than that of HF by 24.86% and 24.52% at seedling and flowering stages; the content of propionic acid was significantly higher than that of HF by 7.36%, 9.44% and23.47% in each growth period, respectively; and tartaric acid acetic acid also showed the same trend at flowering and maturity stages. In summary, tartary buckwheat may regulate the nutrient availability of rhizosphere soil through the secretion of organic acids in the root system to cope with the low nitrogen stress environment. For the cultivation of tartary buckwheat on poor soil should consider the differences cultivaring barren resistance varieties to increase efficiency in the future.

## Introduction

Nitrogen is one of the basic nutrients required for plant growth, which affects crop growth, yield and quality [1]. The lack of nitrogen can lead to slow plant growth, decrease yields, and reduce quality. Therefore, the soil must contain enough nitrogen to meet the normal growth of plants [2]. But nitrogen is a limiting factor in many arable soils in China. For example, the major characteristics of Loess Plateau soils are nitrogen deficiency, low phosphorus and sufficient potassium so that a large amount of nitrogen fertilizer is applied to ensure yield during planting [3]. However, nitrogen fertilizers have lower utilization efficiency, the loss of a large amount of nitrogen not only causes huge waste, but also causes problems such as water bodies and environmental pollution [3]. Therefore, focusing on the overall objective of building a green, efficient and sustainable agricultural production system, the selection of crops with low nitrogen tolerance can not only ensure the normal growth of crops, but also control the amount of nitrogen fertilizer and improve the utilization efficiency of nitrogen and it has significance for improving the ecological environment [4].

Many scholars have conducted research on the low-nitrogen-resistant characteristics of large crops such as wheat and corn [5], but there are few researches on small grains such as tartary buckwheat. Most of the buckwheat is grown in the alpine region, which has the characteristics of short fertility cycle, strong adaptability, drought tolerance and barrenness, and it has extremely high edible value and medicinal value, which is an important disaster relief and pioneer crops [6]. The research on tartary buckwheat mainly focuses on the edible medicinal value, germplasm screening [7]and cultivation techniques, but few studies on the underground parts of tartary buckwheat, especially the response mechanism of tartary buckwheat to low-nitrogen stress are rarely reported.

Studies have shown that after suffering stress from soil nutrient, plant roots first feel the stress and react quickly to adapt to the stress environment through a series changes of root physiological, especially some stress conditions can induce the plant roots to secrete large amounts of organic acids, which is a common active adaptive response [8]. The articles on organic acids secreted by root system under, low nitrogen stress mainly focus on oats, rice and other crops, and also have different results. With the growth of oats, the total amount of organic acids gradually decreased in root exudates, and the total amount of organic acids secreted by the oat roots under nitrogen stress was significantly higher than that of nitrogen supply [9]. Low nitrogen or nitrogen deficiency to accelerate root aging, significantly decrease root secretion capacity and various organic acids of different degrees in rice maturity [10]. Other studies show that there are significant differences in the types and quantities of crop root exudates of the same type of different genotypes [11], and there are different responses to environmental stresses among different cultivars of the same crop [12]. After the determination of various indicators of different genotypes of tartary buckwheat, studies found that the resistance to barrenness of Diqing buckwheat was strong, while the resistance to barrenness of HeiFeng 1 was weak [7].

Therefore, this paper take two different barren tartary buckwheat Diqing and Heifeng 1 as material to compare the differences of root exudation of organic acids in different tartary buckwheat under different nitrogen treatments by pot experiments, in order to explore the response mechanism of buckwheat with different barren conditions to low-nitrogen stress, so as to increase the understanding of tartary buckwheat resistance barren characteristics. It can provide a scientific basis for farmland nitrogen fertilizer optimization management in barren areas of the Loess Plateau.

## Materials and methods

### Test materials

The typical loess was used for the test soils, and the tested crops were buckwheat varieties with different barrenness: Heifeng 1 (HF) and Diqing (DQ). The seeds of Heifeng 1 were provided by High Latitude Crops Institute to Shanxi Academy of Agriculture Sciences with weak resistance to barrenness [7]. The seeds of Diqing were provided by Agricultural Science Institute of Diqing Tibetan Autonomous Prefecture with strong resistance to barrenness [7].

### Experimental design

This experiment was carried out in the plastic greenhouse in Shanxi Normal University from May to August 2017. The experiment was set up three treatments, each treatment was set four repetitions, Three treatments were CK (no nitrogen treatment), N1 (low nitrogen treatment), N2 (normal nitrogen treatment), and were applied to the same amount of phosphate and potassium fertilizer. Urea (46.4% of nitrogen content) was applied of 80 mg kg^-1^ to low nitrogen treatment and 160 mg kg^-1^ to normal nitrogen treatment, phosphate fertilizer was P_2_O_5_ with 150 mg kg^-1^, and potassium fertilizer was K_2_O with 60 mg kg^-1^. Each treatment was set two levels with two different barren buckwheat varieties: HF and DQ. The pot size is 35cm×35cm full of 10kg loess (The soil total nitrogen content of 0.105g kg^-1^) mixed with the fertilizer based on the treatments. 16 full and uniform seeds without pests and diseases were selected and sowed evenly in each pot after soaking in deionized water for 24 hours, 400ml water was used into each pot daily.

### Sample collection

The soil samples were collected in 30d, 60d and 90d after sowing which were represented as Seedling stage(S), Flowering stage(F) and Maturity stage(M) of buckwheat. The soil sample was dried for one week after miscellaneous, grinding, screen and other steps for the determination of soil physical and chemical properties. A portion of the fresh soil was stored in 4°C climate chamber for the determination of NH_4_^+^ and NO_3_^−^, and another portion of fresh soil was stored in -20°C refrigerator for the determination of organic acids.

### Soil properties and Organic acids

Soil Moisture was determined by drying at 105 °C for 12 h. Soil pH was determined in 1:2.5 (soil: water) solution (w/v). Determination of NH_4_^+^: Under basic conditions, NH_4_^+^ reacts with Sodium salicylate and active chlorine to form a colored complex, which is measured at a wavelength of 660 nm [13]. Determination of NO_3_^−^: Under acidic conditions, nitrites react with sulfonamides to produce diazonium compounds, followed by a pink mixture with N-(1-naphthyl)-ethylenediamine dihydrochloride, which is measured at a wavelength of 520 nm [13].

Weigh 2.5g soil sample in 10mL centrifuge tube and add 5mL 0.1% H_3_PO_4_ then, centrifuged at 5000 r min^-1^ speed for 5 min after shock 1 min, passing through the 0.45 um filter in the water phase [14]. Determination of organic acids using HPLC method with Agilent 1290 UHPLC High Performance Liquid Chromatograph (Quaternary Pump, Diode Array Detector, Ultra-efficient autosampler, Temperature intelligent column thermostat). Using CAPCellPAK C18MG4.6mm 250mm, 5um, settings mobile phase for 98:2 ( 0.1% H_3_PO_4_: acetonitrile) (V/V), controlling pH for 2.10, the column temperature for 35°C, the flow rate for 1ml min^-1^, the injection volume for 20ul and the detector wavelength for 210 nm [14-15].

### Statistics and analysis

Using Excel 2010 to statistics the experimental data, SPSS 22.0 software was used to perform multiple comparisons, T-tests, correlation analysis and split plot analysis on the statistical results of the data, and the origin 8.0 software was used for mapping.

## Results

### Effect of buckwheat cultivars and nitrogen treatments on soil moisture

Different N treatments have an impact on soil moisture of HF and DQ (Table 1). In the seedling stage, the soil moisture of HF and DQ under Low N treatment decreased by 24.2% and 14.32%, respectively when compared with the normal N treatment. In the flowering stage, the water consumption of HF under low N treatment was higher than that of normal N treatment, while DQ had no significant difference. In the maturity stage, HF had no significant difference, however the water consumption of DQ is larger than normal N treatment. Under low N treatment, the soil water content of HF was 33.64% higher than DQ in seedling stage, but no significant difference in flowering and maturity (Table 1). This indicates that the water consumption of DQ is significantly higher than that of HF under the low N treatment at seedling stage.

**Table 1.** Soil physical and chemical properties under different tartary buckwheat and nitrogen treatments

| Growth<br>period | Project | Heifeng 1 |  |  | Diqing |  |  |
| --- | --- | --- | --- | --- | --- | --- | --- |
|  |  | CK | N1 | N2 | CK | N1 | N2 |
|  | pH | 7.82±0.03Aa | 7.81±0.07Aa | 7.77±0.05Aa | 7.71±0.05Ba | 7.7±0.04Ba | 7.63±0.04Bb |
| Seedling | SM | 27.3±2.33Aa | 19.8±1.88Ab | 26.1±1.87Aa | 16.2±1.16Bab | 14.8±1.47Bb | 17.3±1.76Ba |
| stage | NH<sub>4</sub><sup>+</sup> | 1.359±0.14Bb | 2.978±0.27Aa | 3.225±0.12Aa | 2.284±0.21Ac | 2.822±0.10Ab | 3.367±0.19Aa |
|  | NO<sub>3</sub><sup>-</sup> | 1.059±0.18Ac | 1.477±0.09Bb | 2.131±0.08Aa | 1.108±0.13Ac | 2.203±0.07Aa | 1.928±0.12Bb |
|  | pH | 7.84±0.05Ab | 8.00±0.02Aa | 8.03±0.05Aa | 7.57±0.03Ba | 7.32±0.07Bb | 7.17±0.07Bc |
| Flowering | SM | 29.1±1.48Aa | 13.4±2.19Ac | 19.9±2.40Ab | 17.1±1.73Ba | 11.2±2.24Ab | 12.5±1.63Bb |
| stage | NH<sub>4</sub><sup>+</sup> | 3.448±0.30Ab | 4.062±0.26Aa | 4.395±0.19Aa | 3.225±0.19Ac | 3.681±0.19Ab | 4.018±0.16Ba |
|  | NO<sub>3</sub><sup>-</sup> | 1.581±0.13Bc | 3.007±0.17Ba | 2.719±0.13Bb | 2.237±0.17Ac | 3.395±0.23Aa | 2.927±0.11Ab |
|  | pH | 8.35±0.02Aa | 8.30±0.03Ab | 8.20±0.01Ac | 8.31±0.06Aa | 8.26±0.04Aa | 8.23±0.05Aa |
| Maturity | SM | 24.2±1.79Aa | 13.4±2.14Ab | 13.4±2.64Ab | 25.5±2.98Aa | 15.4±2.15Ab | 11.5±2.12Ac |
| stage | NH<sub>4</sub><sup>+</sup> | 1.544±0.17Ab | 2.788±0.13Aa | 2.594±0.11Aa | 1.624±0.19Ac | 2.561±0.11Ba | 2.252±0.16Bb |
|  | NO<sub>3</sub><sup>-</sup> | 0.987±0.18Ac | 1.823±0.10Bb | 2.392±0.13Aa | 1.051±0.12Ac | 2.125±0.10Ab | 2.330±0.14Aa |

The values in the table are mean ± standard deviation. Capital letters indicate significant differences in the level of 0.05 for different cultivars of the same N treatment. Lowercase letters indicate significant differences in the level of 0.05 for different N treatment of the same cultivars.

### Effect of buckwheat cultivars and nitrogen treatments on soil pH

Under low N treatment, the soil pH values of DQ were 0.95% and 2.16% higher than those of normal N treatment in seedling and flowering, respectively. While there was no significant difference in HF (Table 1). At maturity, there was no significant difference in DQ, however the soil pH of HF was 1.25% higher than that of normal N treatment. The effects of different cultivars on soil pH mainly concentrated in the seedling stage and the flowering stage, the soil pH value of DQ was lower than that of HF by 1.44% and 8.44% Under low N treatment (Table 1).

### Effect of buckwheat cultivars and nitrogen treatments on soil NH_4_^+^

There was no significant difference in NH_4_^+^ content of HF soil between N1 and N2 at different growth stages, but they all showed significant differences with CK. They were 1.192 times and 1.373 times higher than CK at seedling stage, and 17.8% and 27.5% higher than CK at flowering stage, and 80.6% and 68% higher than CK at maturity stage, respectively (Table 1). The content of NH_4_^+^ in DQ soil showed significant differences under different nitrogen treatments during growth period, N1 was lower than N2 of 16.2% and 8.39% at seedling and flowering stage, respectively, and N1 was higher than N2 of 13.72% at maturity stage (Table 1). Under low N treatment, the content of NH_4_^+^ in DQ soil had no significant difference with HF at seedling and flowering stage, but was lower than that of HF 8.2% at maturity stage (Table 1).

### Effect of buckwheat cultivars and nitrogen treatments on soil NO_3_^−^

Under N1 treatment, the content of NO_3_^−^ in HF soil was lower than N2 of 30.7%, but higher than N2 of 14.3% in DQ soil at seedling stage. the content of NO_3_^−^ in HF and DQ soils were higher than N2 10.6% and 16% respectively at flowering stage, and were lower than N2 23.8% and 8.8% respectively at maturity stage (Table 1). Under low N treatment, the content of NO_3_^−^ in DQ soil was significantly higher than that of HF 49.2%, 12.9%, and 16.6% in each growth period, respectively (Table 1).

### Effect of buckwheat cultivars and nitrogen treatments on the types and contents of soil organic acids

Split plot analysis [16] showed that nitrogen treatment had a significant effect on the organic acid content in the soil of the buckwheat, but the cultivar and interaction had no significant effect on these organic acids (Table 2).

**Table 2.** Interactions between cultivars and nitrogen treatment on soil organic acids

| Project | Significance of difference |  |  |
| --- | --- | --- | --- |
|  | Variety | Nitrogen treatment | Interaction |
| Malonic acid | <0.05 | <0.01 | ns |
| Propionic acid | ns | <0.05 | ns |
| Oxalic acid | ns | <0.01 | ns |
| Tartaric acid | ns | <0.01 | ns |
| Acetic acid | ns | <0.01 | ns |

ns indicate no significant correlation. <0.01 indicate significant differences in the level of 0.01.
<0.05 indicate significant differences in the level of 0.05.

### Types and contents of organic acids in the soil of HF under different nitrogen treatments

In the seedling stage, more propionic acid, oxalic acid and less malonic acid were detected in the soil under low N treatment, but no tartaric acid and acetic acid were detected. At flowering stage, compared to normal N treatment, the roots of buckwheat secrete more propionic acid and oxalic acid, but the secretion of malonic acid, tartaric acid, and acetic acid are reduced by 43.83%, 26.53%, and 23.48% respectively under low N treatment. At maturity stage, the contents of malonic acid, propionic acid, oxalic acid, tartaric acid and acetic acid under low N treatment were lower than that of normal N treatment by 13.84%, 18.56%, 22.84%, 33.40% and 24.19% (Fig 1).

### Types and Contents of Organic Acids in the soil of DQ under Different Nitrogen Treatments

The types and contents of organic acids in the soil of DQ were different under different nitrogen treatment (Fig 2). Compared with normal N treatment, 2.84 times more oxalic acid, 31.54% propionic acid, less 50.87% malonic acid was detected and no acetic acid was detected under low N treatment in the seedling stage. At flowering stage, the roots of buckwheat secrete more oxalic acid and propionic acid in low N treatment, but compared with the normal N treatment, the secretion of malonic acid, tartaric acid and acetic acids decreased by 53.92%, 13.81% and 18.24% respectively under low N treatment. At maturity stage, compared with the normal N treatment, the secretion of buckwheat roots decreased of 23.02% malonic acid, 28.92% oxalic acid, 21% tartaric acid and 24.76% acetic acid in low N treatmnt, but no significant difference was found in propionic acid. In conclusion, the changes of single organic acids content of tartary buckwheat cultivars are different under low N treatments, and it have different performance in the growth period.

### Comparison of organic acids content secreted by roots of different buckwheat under low nitrogen treatment

Under low N stress, in soil of DQ, the content of malonic acid did not differ from that of HF at seedling and flowering stages, but higher than that of HF by 34.39% at maturity stage; The content of oxalic acid was respectively higher than that of HF by 24.86% and 24.52% at seedling and flowering stages, but lower than that of HF by 24.65% at maturity stage; the content of propionic acid was significantly higher than that of HF by 7.36%, 9.44% and 23.47% in each growth period; tartaric acid and acetic acid also showed the same trend, their contents were higher than that of HF by 56.91% and 59.6% at flowering stage, and higher than that of HF by 24.97% and 18.51% at maturity stage, respectively (Fig 3).

### Principal component analysis between soil organic acids and soil properties

The results of principal component analysis showed that there were differences between two varieties and different nitrogen treatments during different growth periods. In the seedling stage (S), the first axis explained 95.9% of variation, which clearly separate N2 from CK, and the second axis clearly separated the two varieties (HF and DQ). In the flowering stage (F), the first axis explained 96.1% of variation, which clearly separated N2 from the other N treatments (CK, N1), but the difference between the two varieties was not significant. In the maturity stage (M), the first axis explained 99.2% of the variation, which clearly separated the two varieties under N1 treatment, but the difference between two varieties was not significant under other N treatments, and the second axis clearly separates CK from the other N treatments (N1, N2) (Fig 4). Except for oxalic acid, the contents of the other four organic acids in the soil showed significant positive correlations at different growth periods, and were negatively correlated with soil moisture and pH, and positively correlated with NH_4_^+^andNO_3_^−^ (Fig 4).

## Discussion

### Effect of Organic Acids on Soil Environment

As the contact surface of plants and soil, roots constantly absorb nutrients and water from the soil to maintain their growth. The root system may secretes various organic acids in rhizosphere when subjected to nutrient stress, which could affect plant growth and development [17]. In this experiment, the principal component analysis showed that malonic acid, propionic acid, tartaric acid and acetic acid had a significant positive correlation with the water consumption of buckwheat, and the water consumption of two tartary buckwheat cultivars under the low N treatment was higher than that under the normal N treatment. Studies have shown that the crop will increase the growth of the main root to absorb water and nutrients deeper and further away from the root system when faced with stress [5]. Therefore, the plants may pass through a large amount of transpiration, allowing more water to carry nutrients into the body for photosynthesis to maintain growth, and at the same time some anti-stress substances are synthesized and secreted through organic acids under low nitrogen stress. However, DQ absorbed more water under normal nitrogen treatment during the mature period, probably because DQ is still undergoing photosynthesis and vegetative growth during this period, and needs to consume more water and substances [5]. Secondly, the principal component analysis showed that there was a significant negative correlation between several organic acids and the soil pH, due to the organic acids secreted by the root increase the concentration of H^+^ in the soil and result the decrease of the pH [18]. In addition, the analysis showed that there are a positive correlation between the content of several organic acids and the available nitrogen content of soil (NH_4_^+^, NO_3_^−^). Previous studies have shown that organic acids secreted by roots can enhance soil nutrient availability, promote the utilization of potentially effective nutrients in the soil, enhance the soil ability to adapt to nutrient-stressed environments by acidifying, chelating insoluble nutrients and ion exchange and restore effect [19-20]. Therefore, the secretion of organic acids from the roots may accelerate the conversion of urea in the soil and increase the availability of the available nitrogen components. Studies have shown that less phosphorus can induce plant roots to secrete large amounts of organic acids, which is of great importance to promote the utilization of insoluble phosphorus-containing compounds in soil [21]. Organic carbon and nitrogen released into soil by the exudation, of which 90% are reabsorbed by the root, promote the material circulation and energy flow of plant nutrient elements [22]. In addition, the low-molecular-weight organic acids secreted by roots can also provide the necessary carbon source and energy for soil microorganisms [23], which can promote the decomposition and transformation of soil organic matter then release nutrients for plant absorption and utilization which could promote plant growth and development.

### Effect of different nitrogen treatments on organic acids secreted by roots of Tartary buckwheat

For a long time, the coupling of plant carbon and nitrogen metabolism has been a hot issue in plant physiology and plant nutrition research, and organic acids are the bond of carbon-nitrogen metabolism coupling. On the one hand, nitrogen metabolism affects the level of organic acids in plants; on the other hand, the formation and accumulation of organic acids play an important role in nitrogen metabolism [24]. It can be seen from the above studies that different nitrogen treatments have significant effects on several organic acids, but the effects are different. The content of single organic acids in the low nitrogen treatments also increases or decreases. Studies have shown that citric acid can be detected in root exudates of soybeans when the nitrogen concentration is low. Conversely, higher concentrations of nitrogen are undetectable [25]. In this experiment, higher concentrations of oxalic acid and propionic acid were detected in the soils of both buckwheat cultivars under low N treatment at seedling and flowering stages. At the same time, the content of NH_4_^+^ in DQ soil was lower than that of normal N at seedling and flowering stage, while the content of NO_3_^−^ in both buckwheat soils was higher than that of normal N at flowering stage under low N treatment. When subjected to nutrient stress, roots of different crops secrete a large amount of certain organic acids to adapt to the stress, and promote nutrient absorption and plant growth through the effects[8]. Under nutrient and water stress, A large number of citric acid was secreted by the roots of two-year-old Larix olgensis, followed by malic acid, oxalic acid [20]. Oxalic acid and citric acid exuded from SH40 and Balenghaitang were increased largely under iron-deficiency stress, significantly increased oxalic acid and citric acid exudation is probably responsible for the tolerance of SH40and Balenghaitang to iron-deficiency stress [26]. White lupine roots secrete a large amount of citric acid to activate the insoluble phosphate in the soil under phosphorus stress, thereby increasing the phosphorus utilization in the soil [27]. A large amount of organic acids will be secreted by larch roots When subjected to soil nutrient stress, and the various physiological and ecological functions of seedlings will be adversely affected by these organic acids to prevent the seedling cell membrane system from being destroyed by stress conditions and thereby protecting the cell membrane system [28]. In summary, a large number of organic acids may be secreted by the roots of buckwheat to adapt to low N stress. At the same time, these organic acids will promote the conversion of NH_4_^+^ to NO_3_^−^, which will increase the content of NO_3_^−^ in the soil and accelerate the absorption of nitrogen. Whether this judgment is correct or not requires further study.

In this experiment, the contents of several organic acids detected in the soils of two buckwheat cultivars were lower than that of normal N treatment, the content of NH_4_^+^ detected in the DQ soil was higher than that of the normal N treatment, and the content of NO_3_^−^ detected in the two buckwheat soils was lower than that of the normal N treatment under low N treatment at maturity stage, which may be due to that plant life activity is basically completed at maturity, the demand for nutrients is reduced, and the content of various organic acids secreted by the root system is also decreased, simultaneously the nitrogen content in the soil was less at maturity. Therefore, the content of NO_3_^−^ that can be absorbed was less under low N treatment, and then the contents of organic acids secreted by the root system also decreased. At the same time, the promotion of the conversion of NH_4_^+^ to NO_3_^−^ was reduced, resulting in more NH4+ content and less NO_3_^−^ content in the soil. Low N or N deficiency accelerated the root senescence, the root exudation ability was significantly decreased, and the organic acids secreted by the root decreased in varying degrees [10]. In the above study, the pH of DQ soil was higher than that of normal N treatment at seedling stage and flowering stage and the pH of HF soil was higher than that of normal N treatment at maturity under low N treatment. This may be due to the total amount of organic acids secreted by the root of the buckwheat were more under the normal N treatment, resulting in a lower soil pH value. Another study shows that nitrate ion is the most important form of inorganic nitrogen absorbed by plant roots. Once it enters the cell, it will be converted into ammonium ions, which in turn participate in the synthesis of amino acid. Plant cells absorb NO_3_^−^ and synergistically absorb H^+^, plants generally continue to release H^+^ to the outside of their cells through respiration, or release the acidic substance to the environment through secretion in order to maintain intracellular pH balance(Ji et al.,2012). Available NO_3_^−^ is relatively less under nitrogen-deficient conditions, and thus the concentration of organic acids released to the environment is relatively low. Conversely, higher concentrations of organic acids are released to the environment under high nitrogen conditions [29].

### Effects of low nitrogen stress on root exudation of organic acids in different Tartary buckwheat

Nitrogen is the most needed nutrient for crop growth and development and it is also a major limiting factor for crop growth. Low nitrogen has a significant effect on plant growth and fruit quality, but there are obvious genotypic differences in the degree of impact [5]. Low-nitrogen tolerant cultivars may have a stronger response mechanism, which make them less affected by low nitrogen stress. Previous studies of different barren crops under low nitrogen stress mainly focused on the differences of their morphological, physiological and root activities. With the increase of low nitrogen stress, The root length and root thickness of low N resistant maize varieties were increased and reduced respectively, and the root morphology is better, root physiological activity and tolerance to low nitrogen stress is strong in order to maintain a more stable growth [30]. Low nitrogen tolerant maize varieties could maintain high root activity under low N stress, thus the nutrient absorption and transport of root system was promoted and the carbon and nitrogen cycle was well maintained [31]. Low-nitrogen-resistant buckwheat cultivars could maintain higher activities of SOD and POD and produce less MDA under low N stress. Higher root activity, NR activity and protein content making it better able to adapt to low-nitrogen environment [5]. In low phosphorus soils, some plant species can grow normally, but some plants are blocked or even die. *C.ichangensis Swing.* the most resistant to phosphorus deficiency, had the highest total amount of organic acid in rhizosphere soil under phosphorus starvation condition. *C.aurantium L.* with the least amount of organic acids in rhizosphere soil had weaker tolerance to phosphorus stress [32].These previous studies have shown that the ability of different genotypes to adapt to stress is different, and barren varieties are more adaptive under nutrient stress. The differences in the types and contents of organic acids secreted by different tartary buckwheat roots may be indicative of their genotypic differences.In this experiment, the water consumption of DQ was significantly higher than that of HF at seedling stage under low N treatment. The reason may not only low-nitrogen-tolerant cultivars had relatively high chlorophyll content, but also photosynthetic system II reaction center was less damaged in low nitrogen environment which could make it a good adaptation. And then photosynthesis can be carried out by absorbing a large amount of water and some anti-stress substances can be synthesized in low nitrogen environment [5]. Under low nitrogen treatment, the content of NH_4_^+^ in DQ soil was lower than that of HF at maturity, and the content of NO_3_^−^ in DQ soil was higher than that of HF at different growth periods. This may be due to that low-nitrogen-tolerant cultivars have a stronger ability to transform nitrogen under low N stress, and convert it from urea to NH_4_^+^, then they can be rapidly transformed into NO_3_^−^ which is better absorbed by the plants, thus the soils of low-nitrogen-tolerant cultivars have a high content of NO_3_^−^. However, the content of NH_4_^+^ in low-nitrogen-tolerant soils was lower at maturity stage, which may be due to the large amount of NH_4_^+^ converted to NO_3_^−^ [33]. In addition, the soil pH of DQ was lower than that of HF at seedling and flowering stage under low N treatment, probably because DQ secreted more organic acids. It turns out that the contents of several organic acids in DQ soil were higher than those in HF under low N stress. In summary, barren-resistant cultivars may adapt to low N stress by secreting more organic acids. Under low N stress, the difference in the secretion of organic acids between Diqing buckwheat and Heifeng 1 may be one of different barren resistance.

## Conclusions

The organic acids secreted by the roots will decrease the soil pH value, accelerate nitrogen transformation and have an impact on the soil environment. Under low nitrogen treatment, the content of single organic acid in the soil is different, and the plant may regulate the nutrient availability of rhizosphere soil through the secretion of several single organic acids in the root system to cope with the stressful environment. In addition, the ability of different genotypes to adapt to stress is different, and the resistant varieties have stronger adaptability. Diqing buckwheat secretes more organic acids than Heifeng 1 under low nitrogen stress, which may be the reason for its strong resistance. For the cultivation of tartary buckwheat on poor soil should consider the differences, breeding barren resistance varieties to increase efficiency in the future.

## Acknowledgements

Special thanks to the seed resources provided by High Latitude Crops Institute to Shanxi Academy of Agriculture Sciences and Agricultural Science Institute of Diqing Tibetan Autonomous Prefecture.

## Supporting information

**S1_Fig.tif.**
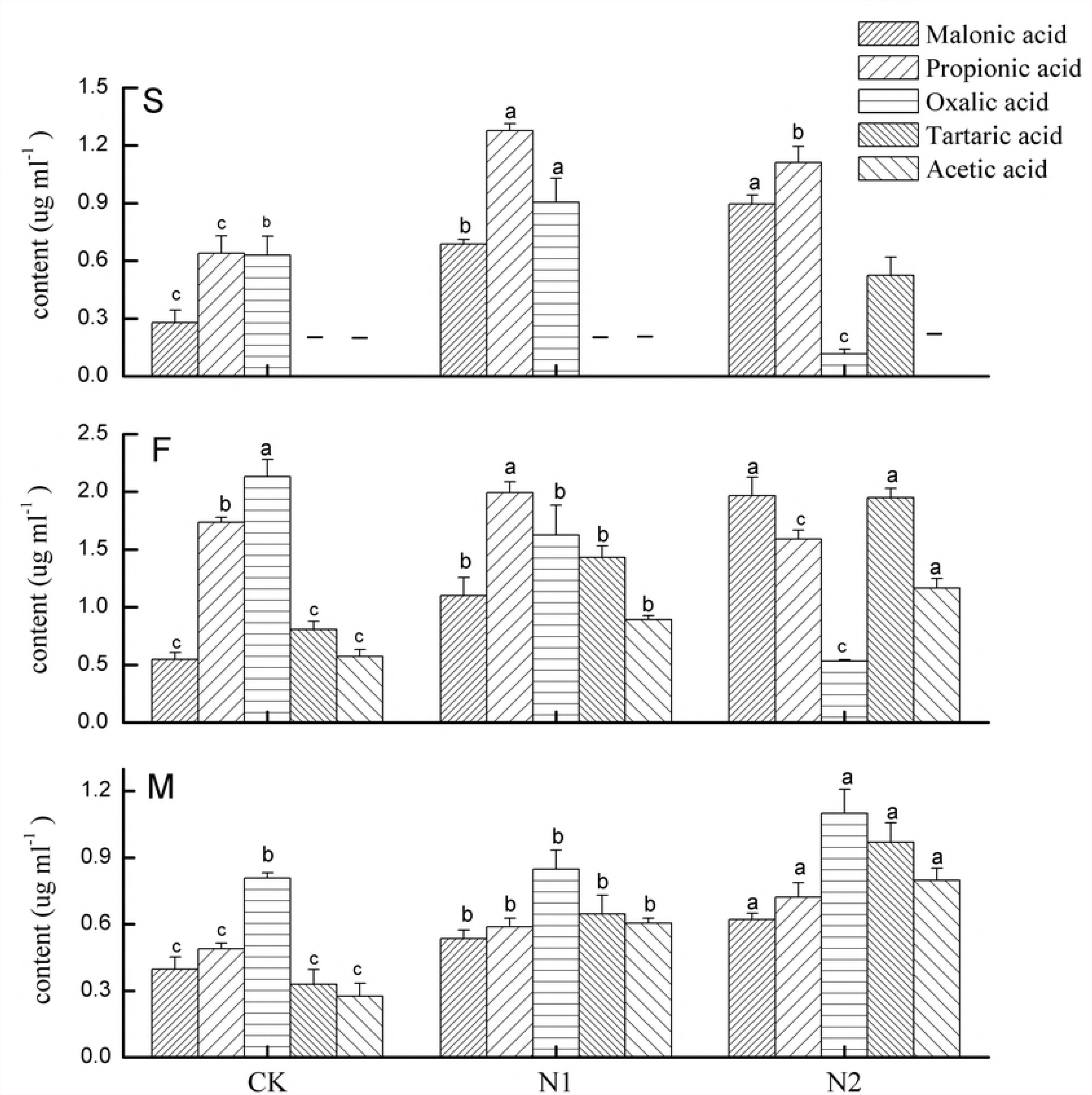
Types and contents of organic acids in HF soil under different nitrogen treatments. S- Seeding stage, F- Flowering stage, M- Maturity stage; CK- No N fertilizer, N1- Low N fertilize, N2- Normal N fertilize. Error bars show the SD. The different lowercase letters indicating significant differences in the level of 0.05 for different N treatments of same organic acid. -Indicates no detected. The same below.

**S2_Fig.tif.**
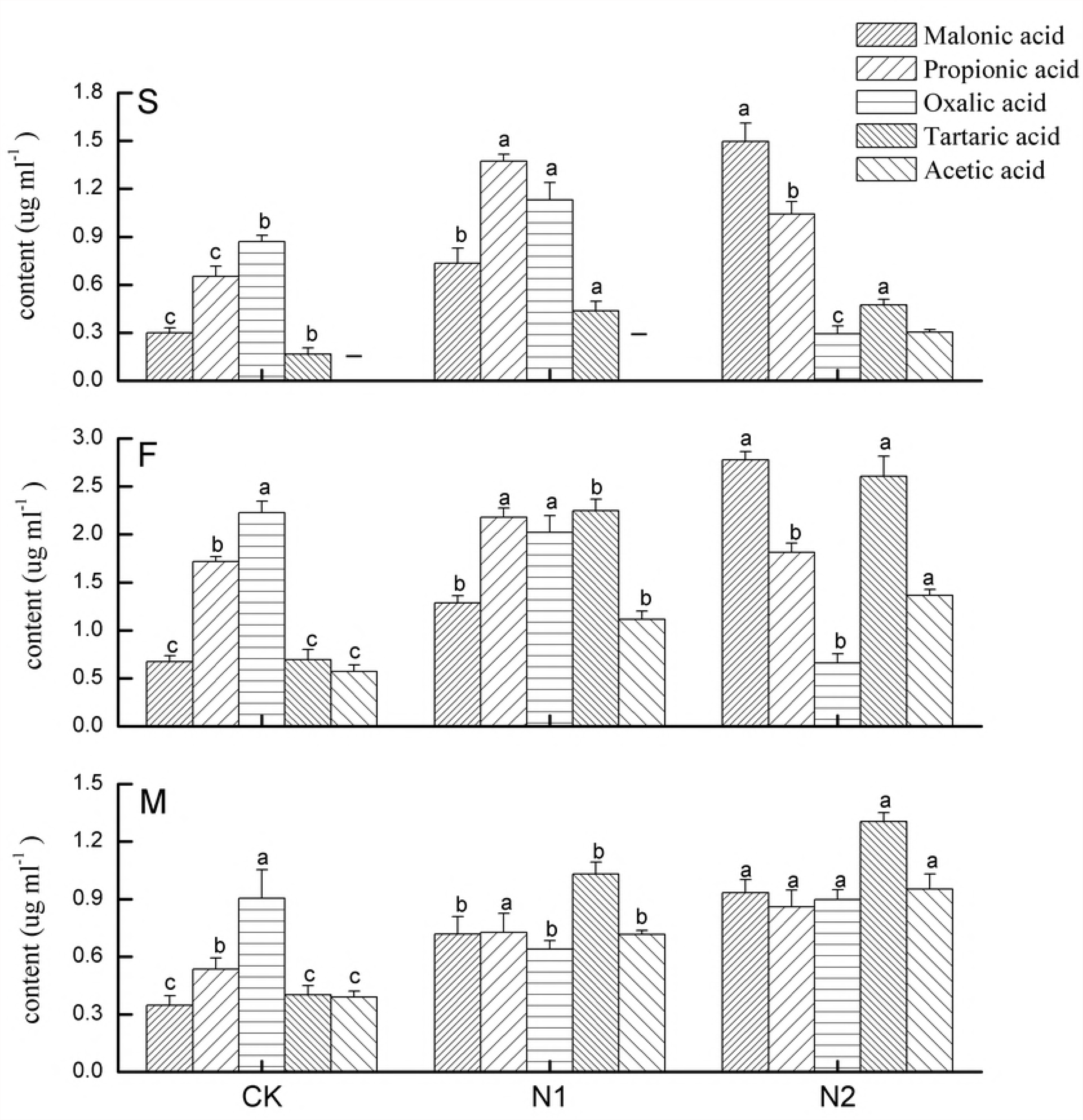
Types and contents of organic acids in DQ soil under different nitrogen treatments. The information refer to the annotation of Fig 1 for explanation.

**S3_Fig.tif.**
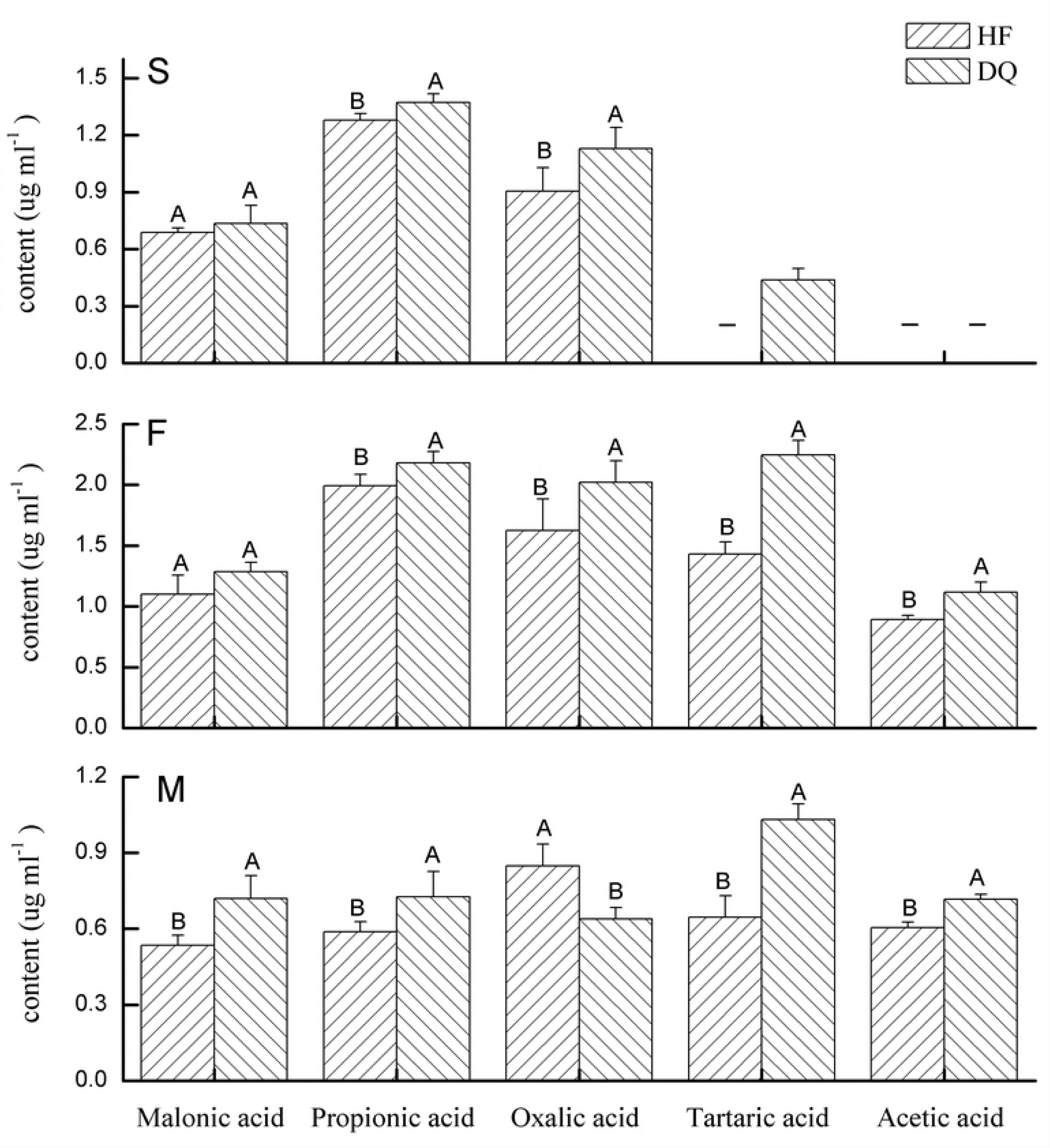
Comparison of the content of organic acids secreted by roots of HF and DQ under low nitrogen treatment. S- Seeding stage, F- Flowering stage, M- Maturity stage; HF- Heifeng 1, DQ- Diqing buckwheat. Error bars show the SD. The capital letters indicate significant differences in probability level of 0.05 for different cultivars of same organic acid. - Indicates no detected.

**S4_Fig.tif.**
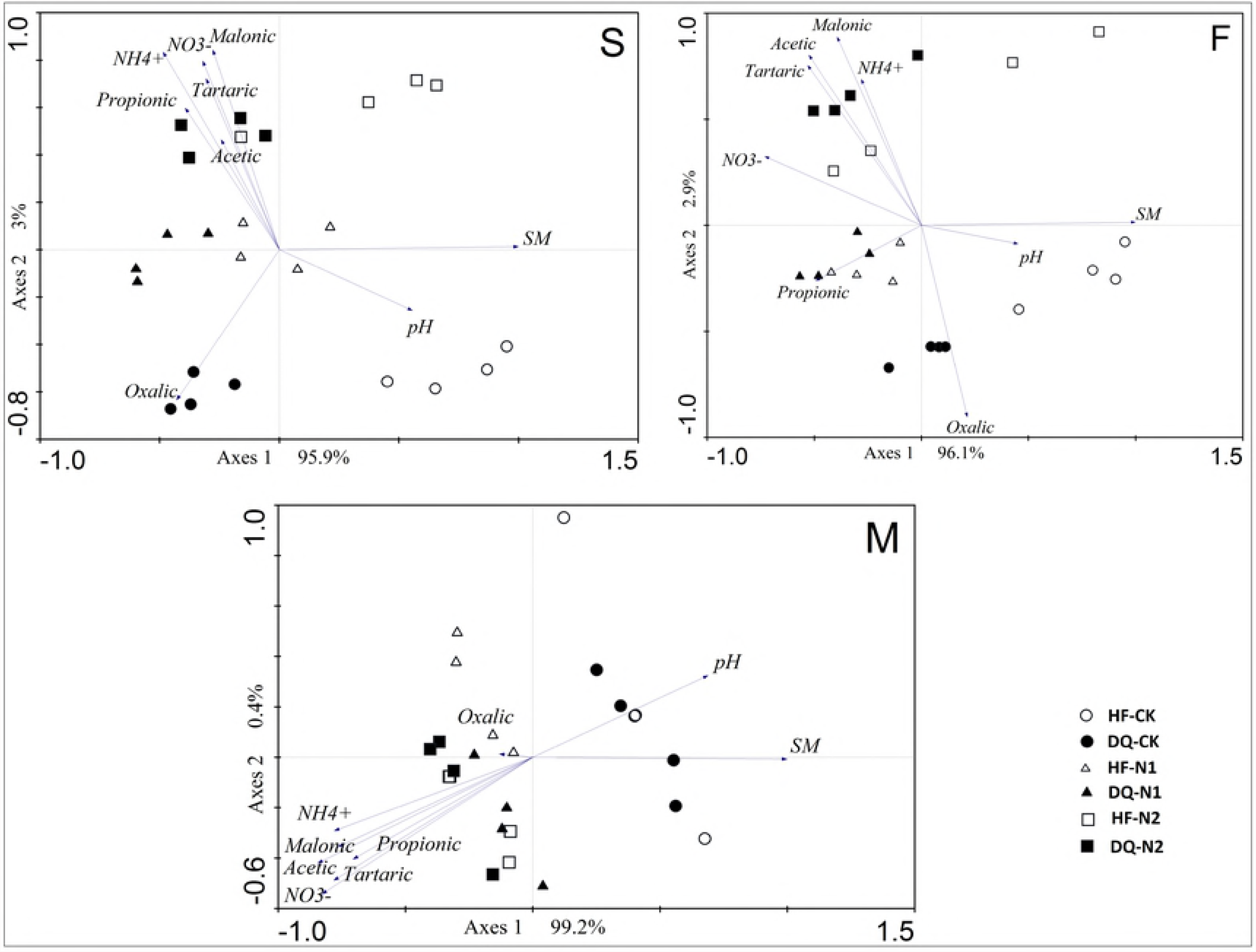
Principal component analysis between soil organic acids and soil properties. S- Seeding stage, F- Flowering stage, M- Maturity stage; HF- Heifeng 1, DQ- Diqing. CK- No N fertilizer, N1- Low N fertilize, N2- Normal N fertilize.

